# A PTN-Rich MSC Subtype Promotes Cartilage Repair in End-Stage Knee Osteoarthritis

**DOI:** 10.64898/2026.11.06.764540

**Authors:** Ziwei Wang, Luxu Yin, Fuchang Zhao, Dan Liu, Jingyu Li, Jing Chu, Zhengtong Zhou, Hui Liu, Rui Fan, Zhenqian Yuan, Yi Zhang, Lingchong Liu, Gongchang Yu, Mengxin Sun, Bin Shi, Jinxiang Han, Yan Zhao

## Abstract

Mesenchymal stem cell (MSC) therapy for end-stage knee osteoarthritis (KOA) shows inconsistent efficacy, and whether MSCs retain cartilage repair capacity in the inflammatory joint remains unclear. Using single-cell RNA sequencing of human Kellgren-Lawrence grade III (G III) and IV (G IV) KOA joint tissues and exogenous MSCs, we found that pleiotrophin (PTN) signalling shifts from MSC-chondrocyte crosstalk to chondrocyte-dominated autocrine signalling, with PTN exhaustion and elevated cluster of differentiation 74 (CD74) in G IV cartilage. We therefore identified and sorted a PTN-rich CD74-low placenta-derived MSC subtype and evaluated it in a murine KOA model and in co-culture with patient-derived chondrocytes. This subtype enhanced cartilage repair, upregulated SOX9, and suppressed MMP13, CD74, IL-6, and IL-1β, potentially through CD74 suppression and reduced NF-κB signaling. These findings provide a basis for functionally defined, stage-matched cell therapy tailored to the pathological microenvironment of end-stage KOA.

**HIGHLIGHTS:**

- Grade IV cartilage shows pleiotrophin exhaustion and elevated CD74-driven inflammation
- A sorted PTN-rich CD74-low placental mesenchymal stem cell subset
- The subset reactivated endogenous chondrocyte repair, raising SOX9, lowering MMP13
- Joint-localized cells persisted 70 days, supporting stage-matched precision therapy

## INTRODUCTION

KOA is a progressive joint disorder that causes pain, functional impairment and structural deterioration[1]. For end-stage KOA, effective non-surgical treatment options remain limited[2]. MSC-based therapy has emerged as a promising disease-modifying approach, however, its therapeutic efficacy remains inconsistent and the mechanisms are poorly understood[2, 3]. This inconsistency may reflect both the complexity of pathological environment in end-stage KOA and the heterogeneity of MSCs, highlighting the need to identify effective MSC-mediated treatment[2, 4]. MSCs have been administered primarily by intra-articular injection, using cells derived from umbilical cord, placenta, bone marrow, adipose tissue and other sources[2]. Clinical studies have primarily assessed pain and joint function, whereas preclinical studies have explored potential paracrine and immunomodulatory effects of MSCs on the joint[2]. While differences in cell source, dose and treatment protocols may contribute to this variability[2, 3, 5], these factors provide limited insight into the specific MSC populations and molecular signals underlying therapeutic benefit. Importantly, MSCs are functionally heterogeneous, and their biological activity may change in disease, with KOA-associated MSCs showing impaired reparative properties and increased pro-inflammatory or matrix-degrading activity[2, 6, 7]. Thus, determining which MSC states and molecular signals are associated with effective tissue repair during disease progression can help clarify the biological basis of therapeutic response and inform strategies to improve MSC-based therapy for late-stage KOA.

Here, we integrated single-cell transcriptomic profiling and intercellular communication analysis of human knee joint tissues from trauma controls and patients with G III and G IV KOA to identify stage-specific molecular signals associated with MSC function during disease progression. We subsequently identified a disease-associated signal and further tested whether a defined MSC subpopulation enriched for this signal could restore chondrocyte reparative capacity in end-stage KOA, and benchmarked this stratified cell-based approach against signal-level modulation alone. This approach may facilitate the identification of potential molecular targets and critical cell subtypes within the pathological microenvironment, thereby informing future strategies to optimise MSC-based therapy for end-stage KOA.

## RESULTS

### Single-cell landscape reveals MSC-derived PTN signaling attenuation accompanied by inflammatory activation during KOA progression

To characterise cellular heterogeneity within the KOA joint microenvironment, we performed single-cell RNA sequencing of cartilage (CA), synovium (SY), and synovial fluid (SF) samples (Supplementary Table S1). Unsupervised clustering and annotation identified 10 major cell populations among 293,499 high-quality cells (Figure 1A). Uniform Manifold Approximation and Projection (UMAP) visualization and celltype composition showed that chondrocytes represented the predominant population, accounting for nearly half of all cells, followed by fibroblasts, macrophages, endothelial cells, endogenous MSCs, and other populations. (Figure 1B)

**Figure 1.**
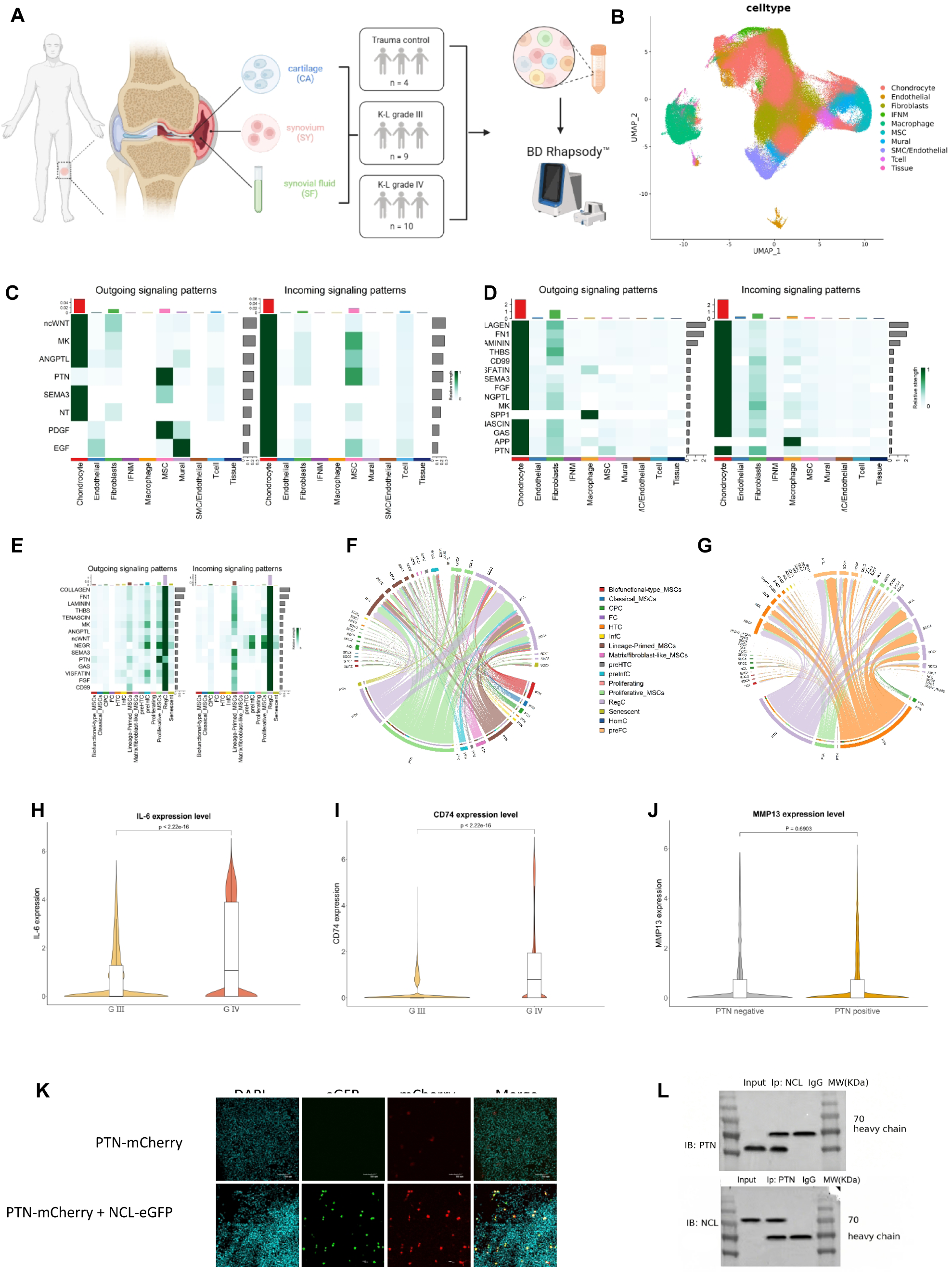
Single-cell transcriptional atlas of human knee joint tissues identifies PTN as a candidate mediator of MSC-chondrocyte crosstalk in KOA. (A) Overview of the study design and sample collection schematic. CA, SY, and SF samples were collected from trauma controls (n = 4) and patients with radiographically confirmed G III (n = 9) and G IV KOA (n = 10). (B) UMAP projection of all 293,499 high-quality cells, colored by 10 major cell types. (C) Outgoing (left panel) and incoming (right panel) signaling patterns across all cell types at G III. (D) Outgoing (left panel) and incoming (right panel) signaling patterns across all cell types at G IV. (E) Outgoing and incoming PTN signaling patterns across subtypes. (F) Subtype-resolved PTN-mediated intercellular communication at G III. At G III, PTN signaling originates predominantly from proliferative MSC and targets RegC. (G) Subtype-resolved PTN-mediated intercellular communication at G IV. At G IV, the MSC-driven axis is diminished, and RegC and HTC emerge as the principal sender populations. Edge width represents communication probability. (H) IL-6 expression across trauma control, G III and G IV groups. IL-6 expression was highest in G IV KOA. (I) CD74 expression in G III and G IV KOA. CD74 expression was significantly increased in G IV compared with G III. (J) MMP13 expression levels in PTN-positive and PTN-negative chondrocytes at G III, with no significant difference between the two groups. (K) Co-localization of PTN-mCherry (red) and NCL-eGFP (green) in co-transfected HEK293T cells. DAPI (cyan) indicates nuclei. Scale bar, 50 µm. (L) Interaction between PTN and NCL was confirmed by reciprocal co-immunoprecipitation. Input, whole-cell lysate; IgG, immunoglobulin G control; IP, immunoprecipitation; IB, immunoblot. Statistical significance is indicated as. \**p*<0.05, \*\**p*<0.01, \*\*\**p*<0.001 and \*\*\*\**p*<0.0001; ns, not significant.

After balancing cell numbers across samples by random downsampling, MSCs were more abundant in CA than in SY and SF. Given that cartilage represents the primary pathological site of KOA, we examined MSC alterations across disease stages in downsampled CA samples.

CellChat analysis revealed extensive intercellular communication changes during KOA progression. Among the identified pathways, PTN signaling represented a major communication axis between MSCs and chondrocytes. MSC-derived PTN signaling was prominent in trauma controls and G III samples, but markedly attenuated in G IV samples (Figure 1C,D). Receptor-level analysis indicated that PTN signaling was primarily mediated through syndecans, including SDC2 and SDC1 (Supplementary Figure S1A). Analysis of matched CA, SY, and SF samples from the same patients further confirmed that PTN expression was consistently higher in CA than in SY and SF at both G III and G IV, supporting the use of CA for subsequent analyses (Figure 1E, Supplementary Figure S1B,C).

### Subtype analysis reveals compositional shifts, Proliferative MSC-RegC PTN signaling diminishment and inflammatory activation in G IV

Given the prominent role of PTN signaling in MSC-chondrocyte communication, we re-clustered chondrocytes and MSCs to define stage-specific cellular states.

Chondrocyte abundance was comparable among trauma controls, G III and G IV CA samples, whereas subtype composition changed substantially during disease progression. Trauma controls were dominated by homeostatic chondrocytes (HomCs), with smaller populations of cycling progenitor cells (CPCs) and pre-hypertrophic chondrocytes (preHTCs) (Supplementary Figure S1D)[8–10]. In G III samples, regulatory chondrocytes (RegCs) became predominant, accompanied by increased cellular heterogeneity. In contrast, G IV cartilage exhibited reduced RegCs and enrichment of hypertrophic chondrocytes (HTCs) and pre-fibrocartilaginous chondrocytes (preFCs), indicating a transition from regulatory to hypertrophic/fibrocartilaginous states during KOA progression.

MSCs displayed distinct stage-dependent alterations, with increased heterogeneity in KOA cartilage. G III samples exhibited enrichment of lineage-primed MSCs and proliferative MSCs, while proliferative MSCs were markedly expanded in G IV cartilage (Supplementary Figure S1E).

We next examined PTN signaling at the MSC and chondrocyte subpopulation levels. Network analysis revealed that proliferative MSCs in G III exhibited strong outgoing PTN signaling despite their limited abundance, acting as the dominant PTN source. RegCs represented the major recipient population, and elevated PTN ligand expression further supported proliferative MSCs as the principal PTN source (Figure 1E). PTN signaling toward RegCs was mediated mainly through PTN-SDC2 and PTN-NCL interactions (Figure 1F). Although SDC2 represented the dominant receptor at the global cell-type level, NCL contribution became apparent after resolving chondrocyte heterogeneity, indicating receptor-level diversity within PTN signaling.

In contrast, PTN-mediated communication was substantially rewired in G IV KOA (Figure 1G). Despite continued expansion of proliferative MSCs, their outgoing PTN signaling was almost completely lost compared with G III, and they were no longer the dominant PTN sender. Instead, RegCs and HTCs emerged as major PTN sources, indicating a transition from MSC-driven to chondrocyte-dominated PTN communication in end-stage KOA. RegC-derived PTN targeted multiple chondrocyte populations, including RegCs, HTCs, and preFCs, further demonstrating extensive remodelling of PTN-mediated communication.

Consistent with the inflammatory phenotype of HTCs, IL-6 expression in CA was significantly increased in G IV versus G III (Figure 1H), indicating a more inflammatory microenvironment in advanced KOA. CD74 expression was also markedly elevated in G IV cartilage (Figure 1I). Given the shift from MSC-derived to chondrocyte-dominated PTN signaling and conflicting reports regarding PTN-mediated matrix degradation, we compared MMP13 expression between PTN-positive and PTN-negative chondrocyte subsets. No significant difference was observed, suggesting that endogenous chondrocyte-derived PTN is not directly associated with MMP13-driven matrix degradation in KOA cartilage (Figure 1J).

To further investigate PTN-receptor interactions, fluorescence co-localization analysis demonstrated partial overlap of PTN-mCherry with NCL-eGFP and SDC2-eGFP in HEK-293T cells (Figure 1K, Supplementary Figure S1F). Co-immunoprecipitation further confirmed physical interactions between PTN and both SDC2 and NCL (Figure 1L, Supplementary Figure S1G). Together, these findings support PTN interactions with SDC2 and NCL at the protein level.

### Single-cell profiling identifies a PTN-rich CD74-low Classical_MSC population in placenta-derived MSCs

Given that the therapeutic efficacy of MSCs in KOA varies across tissue sources, with divergentoutcomes reported for bone marrow-, adipose-, and umbilical cord-derived MSCs[11–13], we therefore performed comparative single-cell transcriptomic profiling of MSCs from various tissue sources to identify a molecularly defined MSC population with therapeutic phenotype.

We established a workflow encompassing MSC isolation, culture expansion and single-cell transcriptomic profiling (Figure 2A). Comparative analysis of human placenta-derived MSCs (hPMSCs) and bone marrow-derived MSCs (hBMSCs) revealed distinct cellular compositions and transcriptional states. UMAP showed that hPMSCs were composed of heterogeneous MSC populations, including Classical_MSC, Proliferative_MSC, etc. (Figure 2B) In contrast, hBMSCs contained a prominent macrophage population in addition to multiple MSC states, indicating a more inflammation-associated cellular composition (Figure 2C).

**Figure 2.**
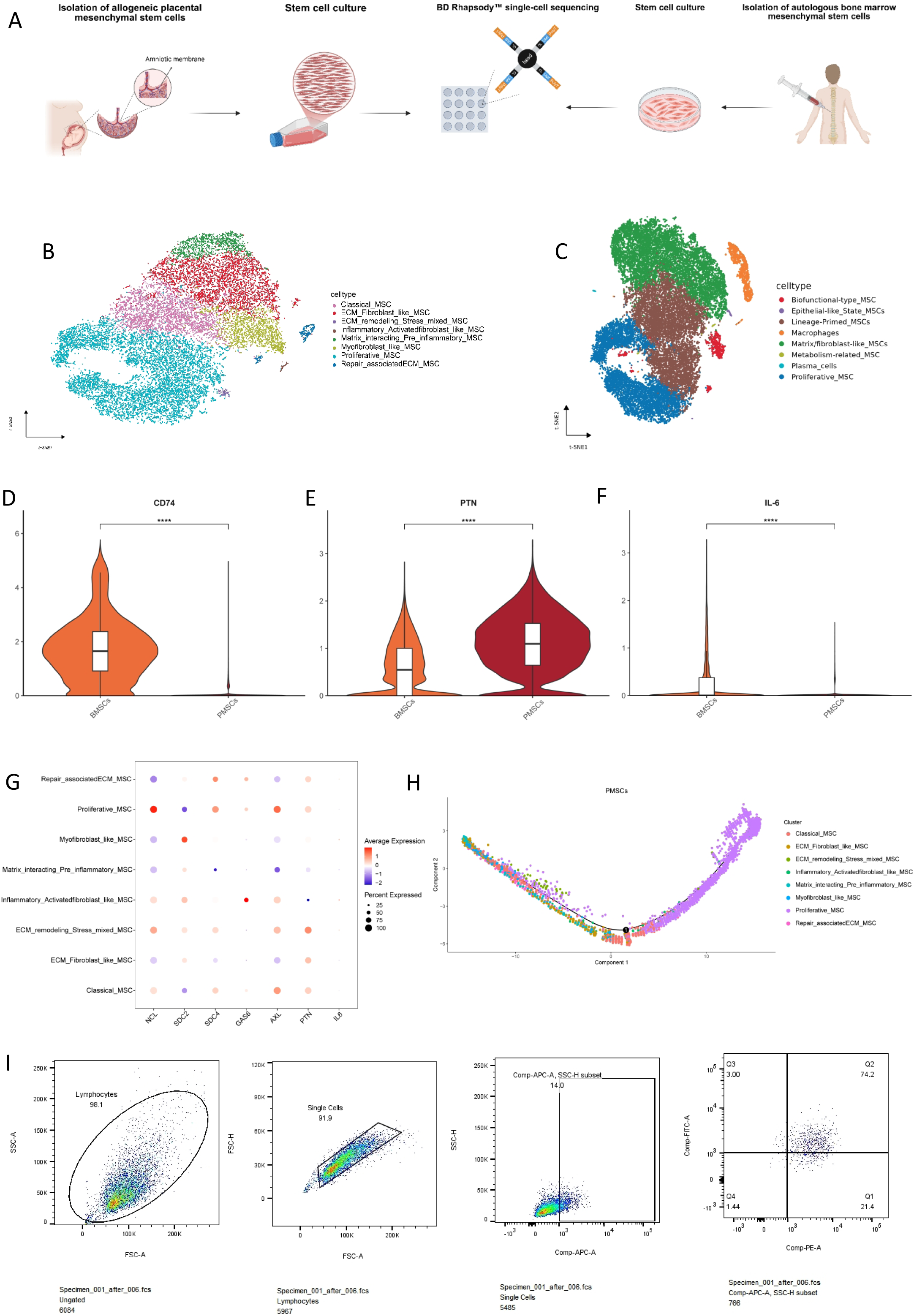
Identification of a PTN-rich CD74-low Classical_MSC subpopulation from exogenous mesenchymal stem cells. (A) Schematic of the isolation, culture and single-cell transcriptomic profiling of hPMSCs and hBMSCs. (B, C) UMAP plots showing the cellular composition and transcriptional heterogeneity of hPMSCs and hBMSCs, respectively. A distinct macrophage population was detected in hBMSCs. (D-F) Violin plots showing the expression of *CD74*, *PTN* and *IL-6* in hBMSCs and hPMSCs. hBMSCs exhibited higher expression of *CD74* and *IL-6*, whereas PTN expression was higher in hPMSCs. (G) Dot plot depicting PTN expression across hPMSCs subpopulations, with the highest expression observed in the Classical_MSC population. (H) Pseudotime trajectory analysis illustrates dynamic functional transitions.

We next asked whether these distinct cellular landscapes were accompanied by differences in molecular features relevant to the inflammatory milieu of end-stage KOA. Notably, CD74 expression was substantially higher in hBMSCs than in hPMSCs, whereas PTN expression was enriched in hPMSCs (Figure 2D,E). Consistent with this pattern, *IL-6* and *MMP13* expression was also higher in hBMSCs than in hPMSCs (Figure 2F, Supplementary Figure S2A), further supporting a more inflammation-associated phenotype in hBMSCs. Given the persistent inflammatory environment of end-stage KOA, the presence of a prominent macrophage population together with higher expression of inflammatory mediators rendered hBMSCs a less favourable candidate for subsequent therapeutic investigation. By comparison, hPMSCs displayed a molecular profile characterised by lower CD74 expression, lower inflammatory signatures and higher PTN expression, providing a rationale for their selection as the MSC source for subsequent studies.

Having established hPMSCs as the preferred MSC source, we next asked whether PTN expression was uniformly distributed across the hPMSC population or preferentially associated with a specific functional state. Subpopulation analysis revealed substantial heterogeneity in PTN expression, with Classical_MSC showing prominent PTN expression compared with other hPMSC states (Figure 2G). Thus, PTN enrichment was not a uniform feature of the hPMSC population but was preferentially associated with a defined MSC state, prompting us to isolate and functionally validate this PTN-rich CD74-low subpopulation as a distinct therapeutic entity rather than relying on unselected bulk hPMSCs.

We therefore investigated the predicted PTN-mediated intercellular communication network within hPMSCs. Ligand-receptor contribution analysis showed that the receptor preference of PTN in hPMSCs was consistent with that in endogenous MSCs (Supplementary Figure S2B). CellChat analysis further delineated PTN-associated communication across hPMSC subpopulations and demonstrated extensive predicted PTN signaling between distinct MSC states. Among these populations, ECM/Fibroblast_like MSC, ECM_remodelling/Stress_mixed MSC and Proliferative_MSC emerged as prominent signaling hubs, contributing substantially to both signal sending and receiving (Supplementary Figure S2C). Together, these findings positioned PTN within a broader communication network across hPMSC states rather than restricted to a subpopulation.

We then determined where the PTN-rich CD74-low Classical_MSC population was positioned within the broader transcriptional landscape of hPMSCs. Pseudotime analysis reconstructed a continuous trajectory extending from the Classical_MSC state towards activated and proliferative_MSC states (Figure 2H). This trajectory suggested that hPMSCs undergo dynamic transcriptional remodelling rather than occupying fixed and independent cellular states. Notably, the PTN-rich CD74-low Classical_MSC population was positioned at an early stage of the inferred trajectory, further distinguishing it from downstream activated and proliferative states. Based on its PTN-enriched phenotype within the hPMSC compartment and the favourable low-CD74 molecular background identified at the source-selection stage, the Classical_MSC was selected as the candidate population for subsequent functional evaluation.

Finally, flow cytometry was used to identify and isolate the Classical_MSC population from cultured hPMSCs. Sequential gating was applied to define the single-cell population and subsequently identify the target MSC compartment (Figure 2I).

Collectively, these analyses established a stepwise strategy for selecting an exogenous MSC population with distinct identities shaped by both tissue origin and molecular characteristics, ultimately identifying a PTN-rich CD74-low Classical_MSC population for subsequent evaluation in experimental KOA.

### A PTN-rich CD74-low Classical_MSC subpopulation enhances functional recovery in KOA

To evaluate the therapeutic potential of the PTN-rich CD74-low Classical_MSC subpopulation in KOA, we established an *in vivo* KOA model and performed intra-articular administration of different therapeutic formulations (Figure 3A). Except for the blank group, all experimental groups underwent bilateral anterior cruciate ligament transection (ACLT) to induce KOA-like joint degeneration. Mice were subsequently randomized into six groups: Blank, Model, Glucosamine, PTN, MSC, and PTN-rich CD74-low MSC (MSC_PTN) groups, enabling systematic comparison of their therapeutic effects on KOA-associated functional impairment.

**Figure 3.**
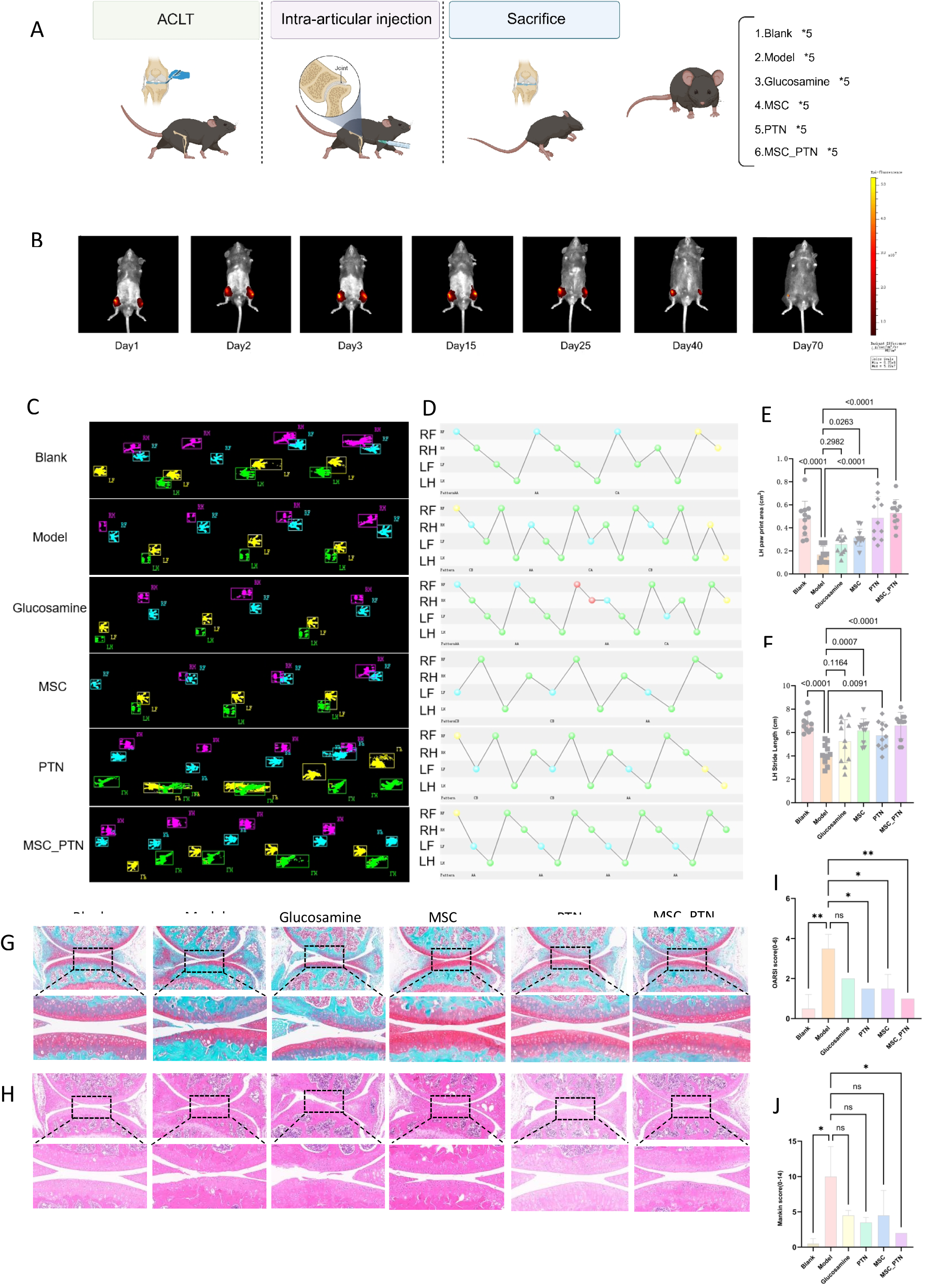
Intra-articular administration of PTN-rich CD74-low MSCs improves joint function and attenuates structural deterioration in experimental KOA. (A) Schematic illustration of the experimental design and treatment groups. Bilateral ACLT was performed to induce KOA in all groups except the Blank group. Mice were randomly assigned to Blank, Model, Glucosamine, PTN, MSC and MSC_PTN groups (n = 5 per group). (B) Representative three-dimensional *in vivo* fluorescence images showing the intra-articular retention of DIR labelled MSCs at days 1, 2, 3, 15, 25, 40 and 70 after injection. (C, D) Representative CatWalk gait recordings illustrating paw-print patterns and gait parameters across the experimental groups. (E, F) Quantification of left hind-limb (LH) paw print area and stride length, respectively. Compared with the Model group, Glucosamine, PTN and MSC treatment produced modest improvements in gait performance, whereas MSC_PTN treatment resulted in a more pronounced improvement in locomotor function. (G) Representative SO/FG-stained sections of the knee joints from each group, with enlarged views of the indicated regions. (H) Representative H&E-stained sections of the knee joints, with corresponding enlarged views. (I, J) Quantitative assessment of cartilage pathology using the knee OARSI and modified Mankin scores, respectively. Data are presented as mean±SD, \**p*<0.05, \*\**p*<0.01, \*\*\**p*<0.001 and \*\*\*\**p*<0.0001; ns, not significant.

We first assessed the intra-articular persistence of transplanted MSCs using longitudinal three-dimensional *in vivo* imaging. hPMSCs showed prolonged and joint-restricted retention after injection, persisting locally for approximately 70 days without evident distant redistribution. (Figure 3B). These findings indicate favorable intra-articular retention of the selected MSC population, supporting prolonged intercellular modulation within the joint microenvironment.

We next evaluated locomotor recovery using the CatWalk gait analysis system. Compared with the Model group, animals receiving glucosamine, PTN, or conventional MSC treatment showed moderate improvements in gait performance (Figure 3C,D). Remarkably, mice treated with the MSC_PTN formulation exhibited substantially superior functional recovery.

Specifically, MSC_PTN-treated mice demonstrated more regular and coordinated gait patterns, accompanied by enhanced hindlimb loading capacity. Quantitative analysis revealed a significant increase in left hind paw print area (*P* < 0.0001) and stride length (*P* < 0.0001) compared with the Model group (Figure 3E,F). Importantly, these improvements were greater than those observed in mice receiving PTN alone or unmodified MSC treatment (Supplementary Figure S3A,B).

We then asked whether functional improvement was accompanied by structural preservation and repair of articular cartilage. Safranin O/Fast Green (SO/FG) staining revealed extensive loss of proteoglycan-rich cartilage matrix and surface irregularity in the Model group, whereas glucosamine, PTN and MSC treatment resulted in varying degrees of attenuation of these pathological changes (Figure 3G). Notably, the MSC_PTN group showed substantially better preservation of cartilage architecture, with a more continuous cartilage surface and stronger SO staining. Consistent with these histological findings, haematoxylin and eosin (H&E) staining demonstrated marked cartilage structural disruption in the Model group, while the MSC_PTN group exhibited a more intact cartilage morphology and reduced tissue deterioration (Figure 3H).

We next used two complementary histological scoring systems, knee Osteoarthritis Research Society International (OARSI) and Mankin scores, to quantitatively evaluate cartilage degeneration and structural damage. Under the knee OARSI score, MSC_PTN achieved a highly significant improvement over the model group (*P* < 0.01), whereas PTN or MSC alone reached only marginal significance (0.01 < *P* < 0.05). Strikingly, under the Mankin score, all other treatments lost significance altogether, leaving MSC_PTN the sole group that remained significantly effective. Regardless of the scoring system, MSC_PTN reproducibly delivered significant efficacy, highlighting its robust and reliable therapeutic effect (Figure 3I,J).

These findings demonstrated that the PTN-rich CD74-low Classical_MSC population provides greater protection against cartilage degeneration and functional impairment than either PTN or unselected MSCs alone. Importantly, these benefits were not confined to a single readout; rather, concurrent improvements across functional, histological, and structural parameters indicate a coordinated, multi-level therapeutic action. Together, these observations support molecular selection as a strategy to refine cell-based therapy for end-stage KOA.

### PTN-rich CD74-low MSCs reverse CD74 elevation and restore joint homeostasis

Having identified PTN as a prominent mediator of MSC-joint crosstalk and a defining feature of the selected therapeutic MSC population, we next sought to delineate how PTN-related signaling interfaces with the molecular landscape of the KOA joint. To this end, we performed in silico gene deletion analysis using a single-cell transcriptomic dataset derived from mouse knee joints. Strikingly, virtual PTN ablation resulted in a substantial upregulation of CD74, which emerged among the most significantly induced genes (Figure 4A). Notably, this effect was not specific to PTN perturbation; virtual deletion of NCL and SDC2, candidate receptors implicated in PTN-mediated signaling, similarly elicited marked CD74 elevation (Figure 4B, Supplementary Figure S4A). These findings directed our subsequent investigation towards CD74 as a potential molecular hub linking PTN-associated signaling to the joint microenvironment.

**Figure 4.**
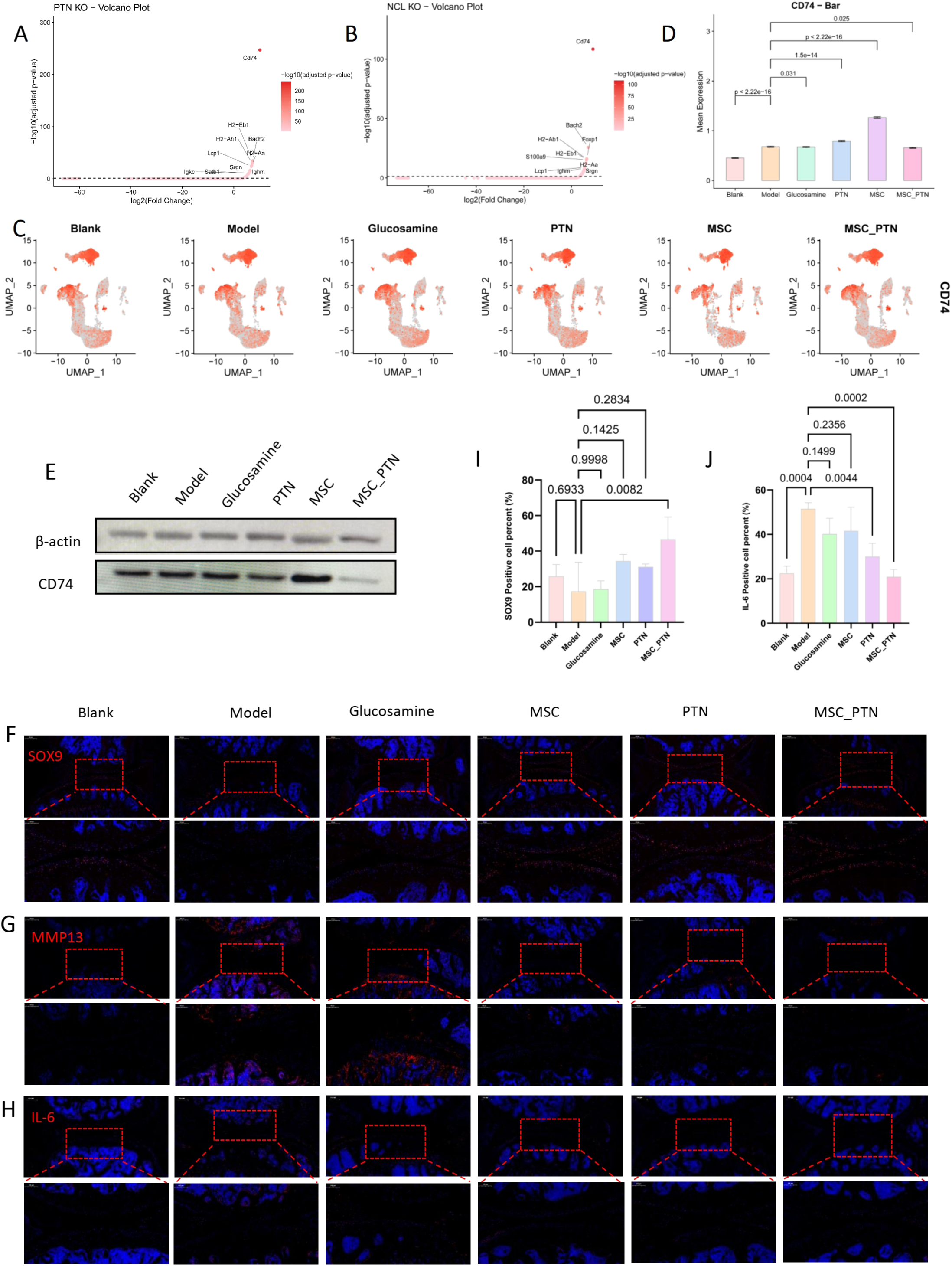
PTN-rich CD74-low MSCs suppress CD74-associated inflammatory and catabolic responses in KOA joints. (A) Volcano plot showing transcriptional changes in mouse knee joint cells following in silico PTN deletion. (B) Volcano plot showing transcriptional changes following in silico deletion of NCL. (C) UMAP feature plots showing the distribution and expression of CD74 across knee joint cells from the blank, model, glucosamine, PTN, MSC and MSC_PTN groups. (D) Quantification of CD74 expression across the six experimental groups. (E) Representative western blot analysis and quantification of CD74 expression in knee joint tissues. (F-H) Representative immunofluorescence images of knee joint sections stained for SOX9 (F), MMP13 (G) and IL-6 (H). (I-K) Quantification of SOX9, MMP13 and IL-6 immunofluorescence signals, respectively. Data are presented as mean±SD, \**p*<0.05, \*\**p*<0.01, \*\*\**p*<0.001 and \*\*\*\**p*<0.0001; ns, not significant.

We next examined whether CD74 expression was modulated by the various therapeutic interventions *in vivo*. Single-cell feature plots revealed distinct CD74 expression patterns across the six experimental groups (Figure 4C). While CD74 remained detectable in KOA joints, its expression was markedly attenuated following treatment, with the lowest signal observed in the MSC_PTN group. Quantitative analysis corroborated this trend (Figure 4D), which was further validated at the protein level by Western blotting, demonstrating a pronounced reduction in CD74 abundance in the MSC_PTN group (Figure 4E). Collectively, these findings indicate that MSC_PTN treatment was accompanied by reduced CD74 expression and was associated with the most favourable molecular and structural outcomes in the KOA joint.

We then asked whether these molecular alterations were accompanied by restoration of cartilage homeostasis. Immunofluorescence analysis revealed a robust increase in SOX9, a master regulator of the chondrocyte anabolic program, in the MSC_PTN group. Conversely, the cartilage-catabolic marker MMP13 and the pro-inflammatory mediator IL-6 were substantially diminished following MSC_PTN treatment (Figure 4F-H). Immunofluorescence quantification confirmed that MSC_PTN significantly increased the number of SOX9-positive cells compared with the Model group (*P*=0.0082), whereas PTN, MSC and glucosamine showed no significant improvement. Conversely, IL-6-positive cells were significantly reduced by PTN (*P*=0.0044) and further reduced by MSC_PTN (*P*=0.0002), with MSC_PTN showing the most pronounced effect. Together, these findings highlighted the substantial effects of MSC_PTN in restoring the anabolic phenotype and suppressing inflammatory activity (Figure 4I,J, Supplementary Figure S4B).

Collectively, these findings were consistently recapitulated across independent experimental paradigms: loss of PTN or NCL was associated with increased CD74 expression, whereas administration of PTN-rich CD74-low MSCs correlated with reduced CD74 expression and improved cartilage-associated molecular features.

### PTN-rich CD74-low MSCs improve cartilage-associated phenotypes in a human KOA-mimicking co-culture system

To further evaluate the effects of PTN-rich CD74-low MSCs in a human KOA microenvironment, we established an *in vitro* co-culture system using chondrocytes from patients with G III or G IV KOA. The PTN-rich CD74-low MSCs were expanded *in vitro* and co-cultured with G III or G IV KOA chondrocytes using Transwell inserts. After 48 hours, cells from the lower chamber were collected for scRNA-seq, together with chondrocytes cultured alone for comparative analysis (Figure 5A).

**Figure 5.**
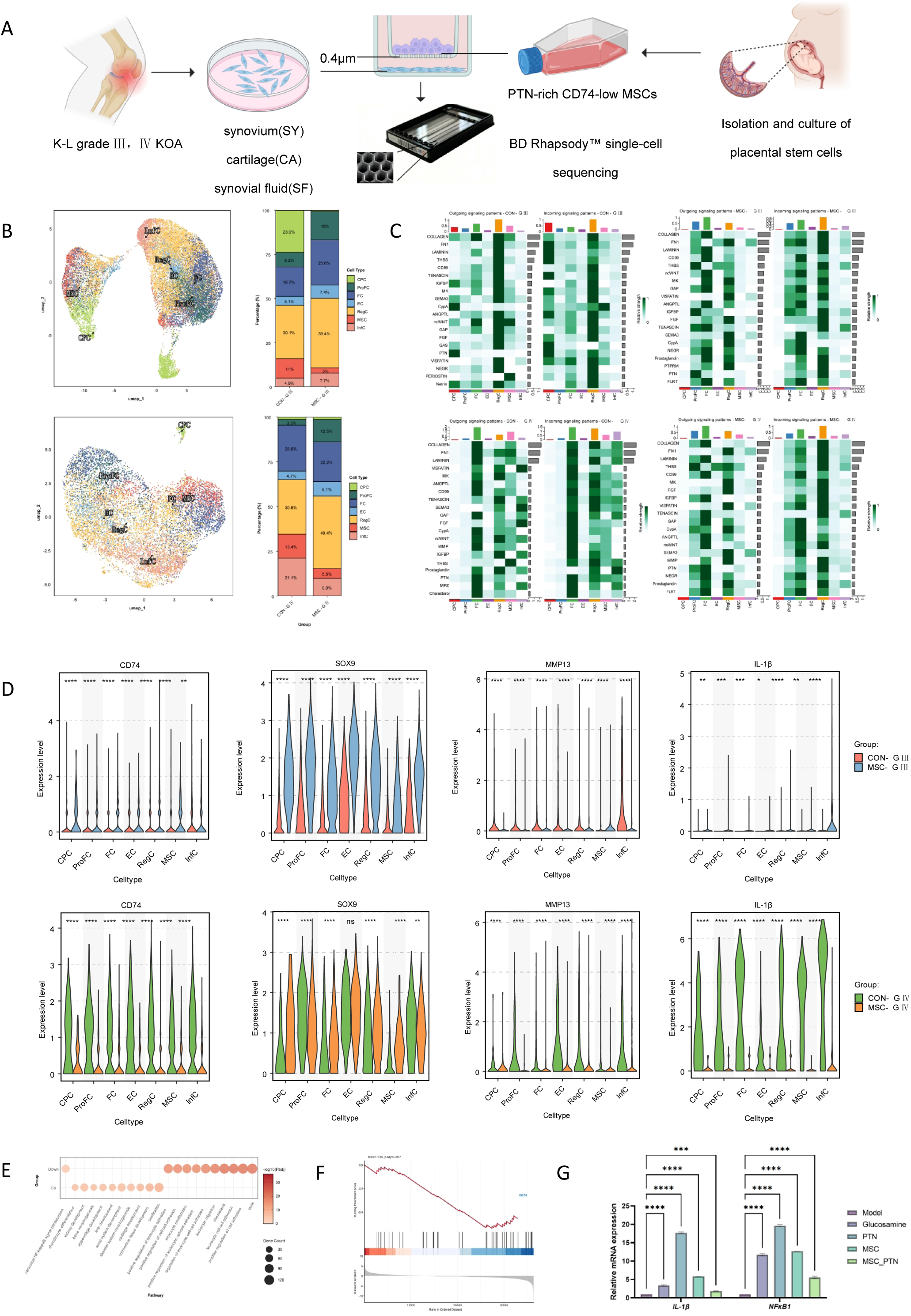
PTN-rich CD74-low MSC suppress inflammation and facilitate cartilage repair by inhibiting the NF-κB pathway via CD74 regulation. (A) Schematic workflow of the co-culture single-cell sequencing procedure. (B) UMAP visualization of cell clustering in G III and G IV chondrocytes. (C) Heatmap of intercellular communication in G III and G IV samples. (D) Violin plots showing the expression levels of PTN, SOX9, MMP13, IL-1β, and CD74 in MSC versus CON groups. (E) Comparative GO enrichment bubble plot of MSC-IV versus CON-IV. (F) GSEA plot comparing MSC-IV versus CON-IV. Data were analyzed using the Wilcoxon rank-sum test. Statistical significance is indicated as follows: \**p* < 0.05, \*\**p* < 0.01, \*\*\**p* < 0.001, and \*\*\*\**p* < 0.0001.

Figure 5B shows UMAP visualisation of cell clustering in G III and G IV samples, with seven distinct subpopulations identified in each group. Before co-culture, CPCs accounted for a higher proportion of cells in G III samples, whereas the proportion of RegCs increased following co-culture. Given the mesenchymal-like progenitor characteristics of CPCs, their greater abundance at G III suggests a higher reparative potential at this stage. In contrast, CPCs were barely detectable in G IV samples before co-culture, indicating a marked reduction of this progenitor population at the advanced stage. Notably, the proportion of RegCs also increased following co-culture. We next examined changes in PTN-associated cell-cell communication following co-culture (Figure 5C). Before co-culture, CPCs represented the predominant source and recipient of PTN signaling in G III samples, whereas RegCs became the dominant recipient following co-culture. In G IV samples, PTN communication was undetectable before co-culture but became evident following co-culture, with fibrocartilage chondrocytes (FCs) representing the predominant source. These findings suggest that PTN-associated communication, which was largely absent in G IV samples, can be restored following co-culture with PTN-rich CD74-low MSCs, accompanied by changes in the chondrocyte subpopulation landscape.

We further assessed cartilage anabolic activity by *SOX9*, cartilage catabolic activity by *MMP13*, and inflammatory status by *CD74* and *IL-1β* expression (Figure 5D). Co-culture induced coordinated changes in anabolic and catabolic markers, with *SOX9* upregulated, particularly in G III samples, and *MMP13* downregulated in both G III and G IV samples. In G IV samples, the inflammatory markers *IL-1β* and *CD74* were also significantly reduced following co-culture. GO analysis further revealed downregulation of the NF-κB signaling pathway after co-culture (Figure 5E). Consistently, GSEA demonstrated a marked reduction in *CD74* expression in G IV samples following co-culture (Figure 5F).

We next asked whether these changes observed in the human KOA-mimicking co-culture system were recapitulated *in vivo*. In the murine KOA model, RT-qPCR analysis showed that MSC_PTN treatment significantly reduced the mRNA levels of *IL-1β* and *NFKB1* compared with the Model group (Figure 5G). The concordant reduction in inflammatory markers in both the human co-culture system and the murine KOA model provides cross-species validation for the anti-inflammatory effects of PTN-rich CD74-low MSCs. Together, these findings indicate that PTN-rich CD74-low MSCs improve cartilage-associated phenotypes and attenuate inflammatory activity across complementary human and murine KOA models.

## DISCUSSION

In this study, by integrating single-cell transcriptomics of human G III and G IV KOA cartilage with *in vitro* and *in vivo* validation, we delineated the molecular transition across disease progression at end-stage and identified an MSC subtype capable of counteracting it. Transition from G III to G IV was characterised by decreased PTN expression, elevated CD74, attenuated reparative communication, and enhanced inflammatory programmes. Against this background, we identified a PTN-rich CD74-low MSC subtype that upregulated SOX9 while reducing MMP13, IL-6, and IL-1β, suppressed CD74-associated NF-κB signaling, localized within joints without off-target diffusion, and persisted *in vivo* for up to 70 days.

Our findings indicate that MSC-mediated reparative activity and PTN signaling patterns were concordant across *in vivo* and *in vitro* experiments. Within the pathological joint environment, G III and G IV KOA exhibited distinct cellular profiles, with MSC-mediated reparative activity progressively declining with disease severity. The increased abundance of MSCs in G III may reflect a compensatory response, whereas their decline in G IV suggests impaired reparative capacity. Consistent with these *in vivo* observations, *in vitro* co-culture experiments revealed stage-dependent responses to PTN-rich CD74-low MSCs. In G III, CPCs acted predominantly as PTN senders, whereas RegCs served as major PTN receivers[14, 15]. By contrast, G IV was characterised by near-complete depletion of CPCs, with fibrocartilaginous cells assuming chondro-reparative functions, indicative of profound regenerative exhaustion. Notably, PTN supplementation partially restored regenerative signaling and increased RegC abundance, suggesting that disease stage may influence the responsiveness to PTN-mediated reparative cues and providing a potential basis for patient stratification and precision treatment. These findings support the rationale for using exogenous MSCs as a therapeutic strategy for KOA, while highlighting that their reparative efficacy may depend on the disease stage and functional state of the administered cells.

The concurrent increase in CD74 and inflammatory features suggests a shift from reparative to inflammatory programmes during progression[16]. Originally characterised as an MHC class II-associated invariant chain, CD74 has emerged as a functional signaling receptor: membrane-associated CD74 binds macrophage migration inhibitory factor (MIF)[17], and activates NF-κB signaling[18, 19]. NF-κB activation drives KOA progression by inducing inflammatory cytokines, matrix metalloproteinases, and catabolic responses[20]. Our study demonstrates aberrant CD74 upregulation in end-stage KOA cartilage with enhanced inflammatory programmes, while PTN supplementation significantly reduces CD74, suppresses NF-κB signaling, and decreases IL-1β, IL-6 and MMP13 with particularly pronounced effects in G IV. These results suggest that CD74 is not merely an inflammatory marker but a critical signaling hub linking inflammation with cartilage degeneration.

These findings also address a long-standing challenge in MSC-based therapy: inconsistent efficacy associated with tissue origin, heterogeneity and functional variability[21]. Although MSC heterogeneity is well recognised, its implications for clinical translation have not been sufficiently considered. In clinical practice, MSC-based therapies are often administered as relatively heterogeneous bulk cell populations, with limited consideration of the functional differences among constituent MSC subpopulations[22]. Consistent with this concern, our single-cell analysis revealed substantial functional heterogeneity within MSCs and identified a PTN-rich, CD74-low subpopulation characterised by enhanced chondroprotective and anti-inflammatory properties. Its prolonged intra-articular retention further supports its therapeutic potential. Our findings suggest that intrinsic heterogeneity may contribute to therapeutic instability by diluting rare potent subpopulations among less effective cells. Neither PTN supplementation nor bulk MSCs reproduced the therapeutic benefit of the selected subpopulation[23], whereas PTN-rich CD74-low MSCs markedly improved efficacy and consistency. Collectively, these findings pinpoint functional heterogeneity as a determinant of MSC efficacy and support a shift from cell quantity-dependent delivery towards functionally defined precision cellular therapy.

Our study does not support the concept of exogenous MSCs as direct substitutes for damaged cartilage. Instead, the semipermeable membrane-based co-culture system, which physically separated MSCs from patient-derived cells, indicates that the effects of MSCs are mediated predominantly through paracrine signaling. Functionally targeted MSCs reshape the intercellular communication landscape and local microenvironment of patient-derived cells, thereby promoting reparative cell expansion, restoring SOX9, reducing MMP13, and attenuating CD74-associated inflammatory programmes. Thus, rather than directly replacing damaged tissue, exogenous MSCs may act as microenvironmental modulators that re-establish conditions in which the patient’s endogenous cells can regain their reparative and homeostatic capacity.

In conclusion, KOA progression is characterised by a shift from PTN-rich CD74-low to PTN-deficient CD74-high states, accompanying exhaustion of cartilage regenerative capacity. Targeting this imbalance through functional MSC selection restored joint homeostasis and improved therapeutic efficacy, presenting a precision cellular strategy for KOA.

### Limitations of the study

This study has several limitations. First, the effects of MSC subpopulations other than the PTN-rich CD74-low subset remain incompletely elucidated. Second, although reduced PTN signaling could induce CD74 over-expression via the NF-κB pathway, with the detailed regulatory mechanisms awaiting further investigation. Moreover, these findings require confirmation in larger-scale studies.

## Supporting information

Supplemental Document S1: Clinical Patient Information Statistics Table and Primer sequences used for RT-qPCR

Supplemental Document S3: BioRender publication license

## RESOURCE AVAILABILITY

### Lead contact

Requests for further information and resources should be directed to and will be fulfilled by the lead contact, Yan Zhao.

### Data availability

Standardized data type data have been deposited at National Genomics Data Center as PRJCA074037 and are publicly available as of the date of publication. Any additional information required to reanalyze the data reported in this paper is available from the lead contact upon request.

## ACKNOWLEDGEMENTS

The authors thank Baheal Pharmaceutical Group Co., Ltd. for providing financial support. We would like to thank Mr. Xu Hao of Wuhan Benagen Technology Co., Ltd. (Wuhan, China), and Mr. Jin Zhiming and Mr. Guo Yichu of OMNIOMICS (Shandong) Biotechnology Co., Ltd. (Jinan, China), for their valuable guidance and discussions. We are also grateful to the Laboratory Animal Center of the Central Hospital Affiliated to Shandong First Medical University (Jinan, Shandong 250013, P.R. China) for providing the mouse gait analysis instrument. Finally, we sincerely thank all the patients who generously donated their samples for this KOA research.

## AUTHOR CONTRIBUTIONS

Ziwei Wang: Writing - review & editing, Formal analysis, Data Curation. Luxu Yin: Writing - review & editing, Investigation, Data Curation. Fuchang Zhao: Writing - review & editing, Validation. Dan Liu: Writing - review & editing, Methodology, Validation, Formal analysis, Investigation, Data Curation, Visualization. Jingyu Li: Writing - review & editing, Methodology, Validation, Formal analysis, Investigation, Data Curation, Visualization. Jing Chu: Writing - review & editing, Methodology, Validation, Formal analysis, Investigation, Data Curation, Visualization. Zhengtong Zhou: Writing - review & editing, Validation. Hui Liu: Writing - review & editing, Validation. Rui Fan: Writing - review & editing, Validation. Zhenqian Yuan: Writing - review & editing, Validation, Visualization. Yi Zhang: Writing - review & editing, Validation. Gongchang Yu: Writing - review & editing, Project administration. Mengxin Sun: Writing - review & editing, Validation. Bin Shi: Writing - review & editing, Project administration. Jinxiang Han: Writing - review & editing, Project administration, Funding acquisition. Yan Zhao: Writing - Original Draft, Conceptualization, Methodology, Resources, Supervision, Project administration, Funding acquisition.

## ETHICS APPROVAL

This study was approved by the Institutional Ethics Committee of the Affiliated Hospital for Neck, Shoulder, Waist and Leg Pain, Shandong First Medical University (Ethics Review Approval No. 202401045) (Jinan, Shandong, China) and the Medical Ethics Committee of the Affiliated Hospital for Neck, Shoulder, Waist and Leg Pain, Shandong First Medical University (approval No.202502027) (Jinan, Shandong, China).

## SUPPLEMENTAL INFORMATION

Document S1. Table S1, Clinical Patient Information Statistics Table; Table S2, Primer sequences used for RT-qPCR.

Document S2. Figure S1, PTN in KOA: expression, subtype distribution, and SDC2 interaction; Figure S2, PTN signaling and receptor-ligand interactions across MSC subpopulations; Figure S3, Identification and validation of SDC2-associated molecular changes and MMP13 expression; Figure S4, Effects of MSC_PTN treatment on hindlimb gait parameters in mice.

Document S3. Confirmation of Publication and Licensing Rights - Open Access.

---

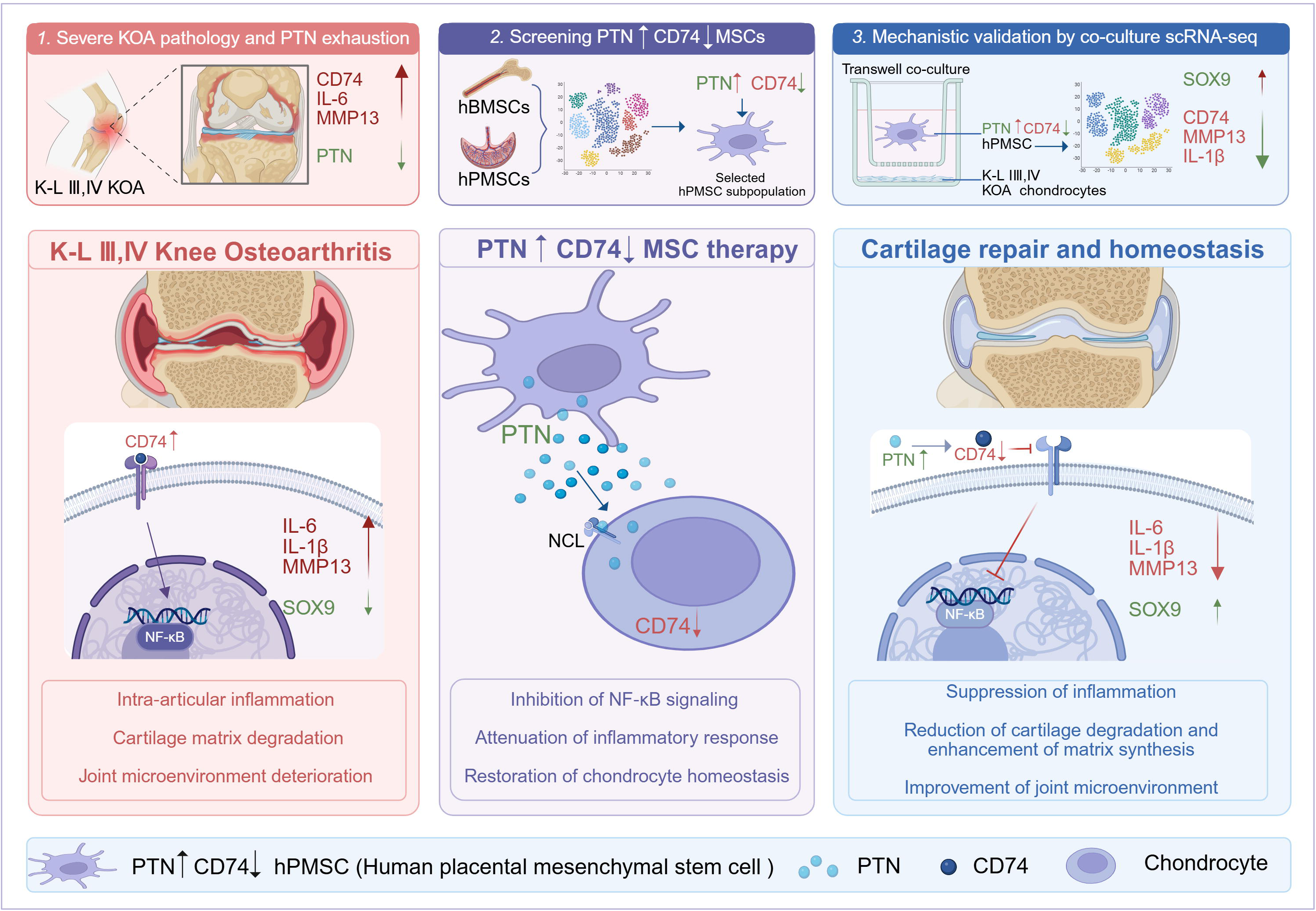

