## Supplemental Document S2: Supplementary figures for "A PTN-Rich MSC Subtype Promotes Cartilage Repair in End-Stage Knee Osteoarthritis"

### Slide 1
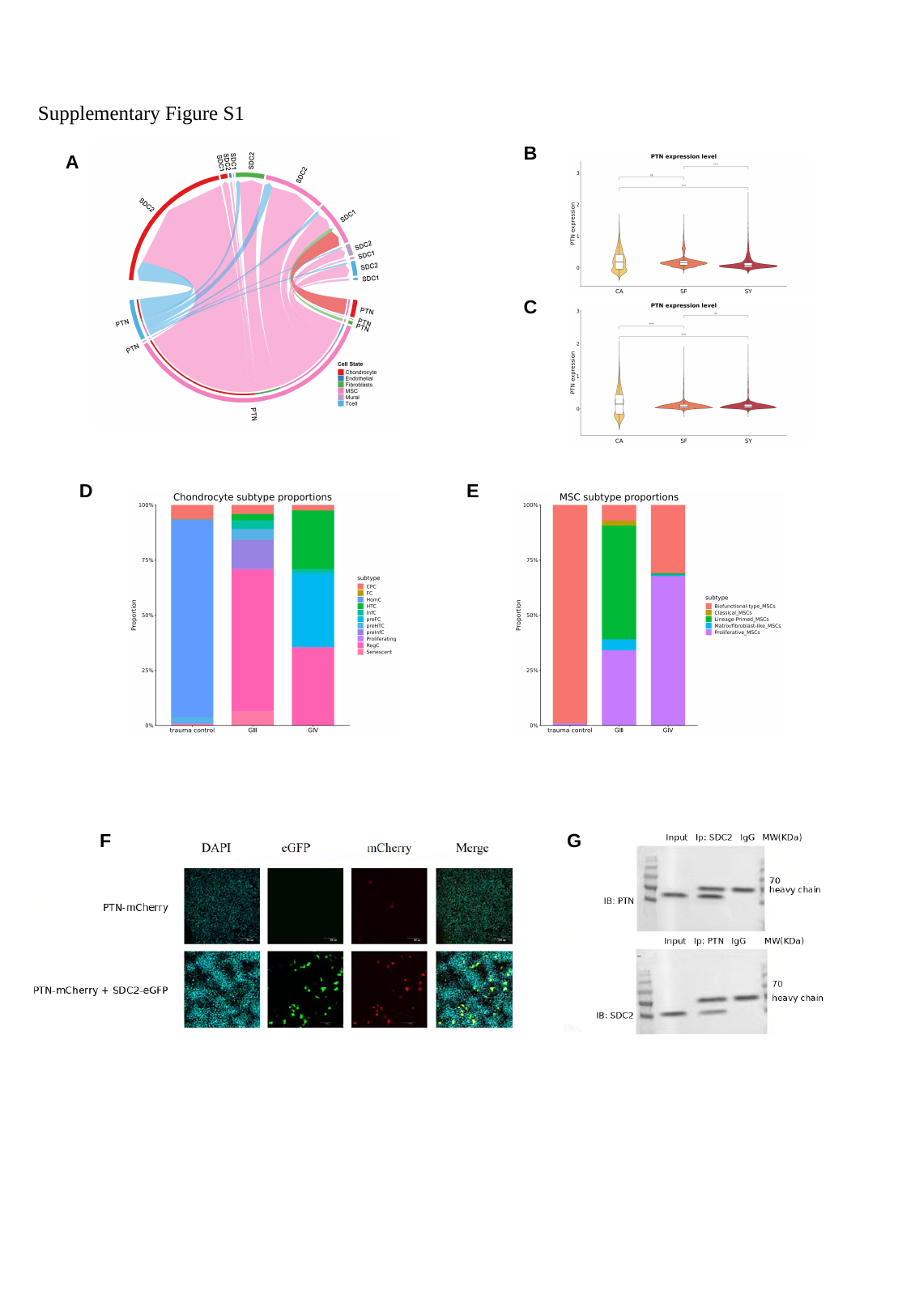

Supplementary Figure S1
B
A
C
D
E
F
G

### Slide 2
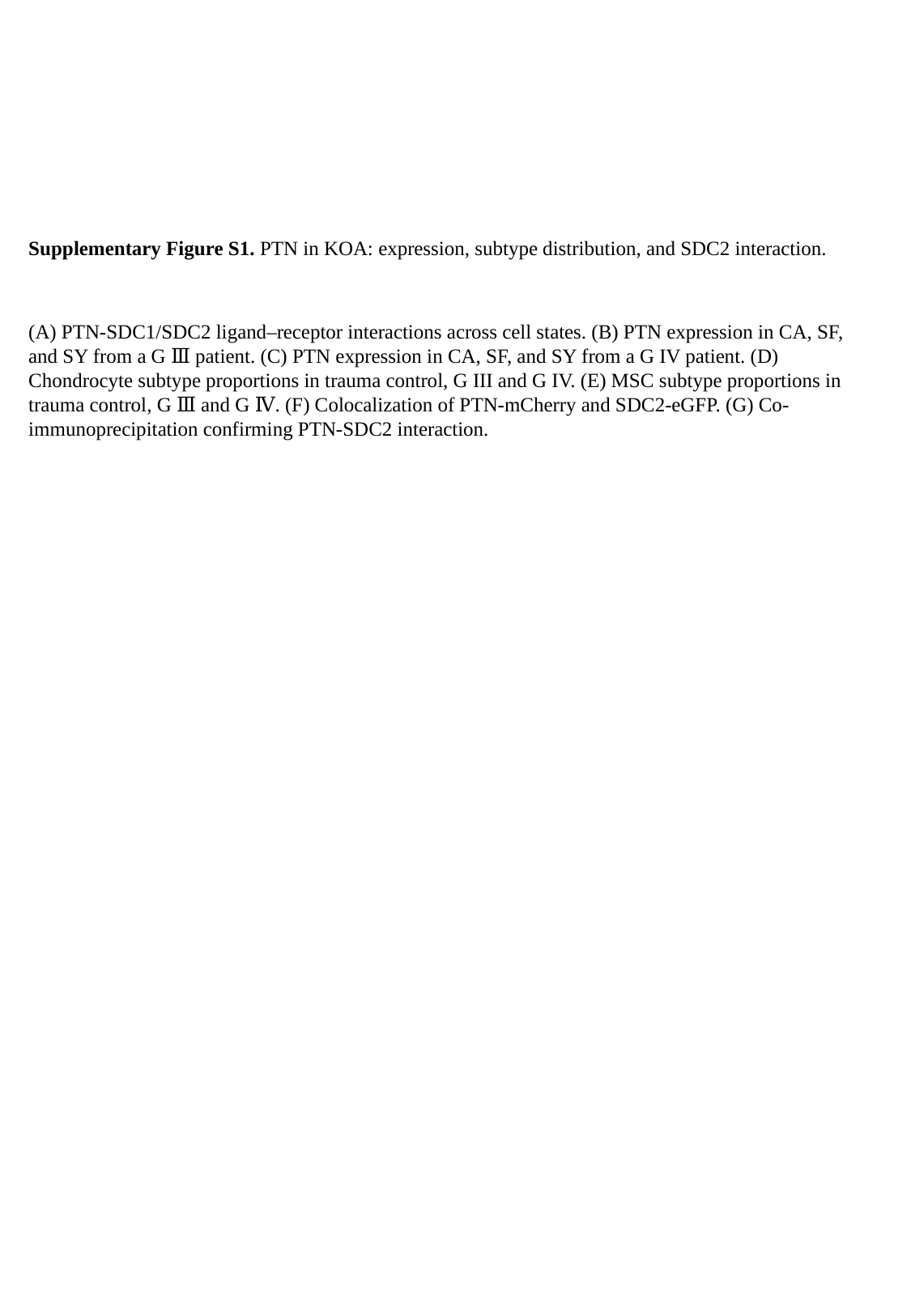

Supplementary Figure S1. PTN in KOA: expression, subtype distribution, and SDC2 interaction.
(A) PTN-SDC1/SDC2 ligand–receptor interactions across cell states. (B) PTN expression in CA, SF, and SY from a G Ⅲ patient. (C) PTN expression in CA, SF, and SY from a G IV patient. (D) Chondrocyte subtype proportions in trauma control, G III and G IV. (E) MSC subtype proportions in trauma control, G Ⅲ and G Ⅳ. (F) Colocalization of PTN-mCherry and SDC2-eGFP. (G) Co-immunoprecipitation confirming PTN-SDC2 interaction.

### Slide 3
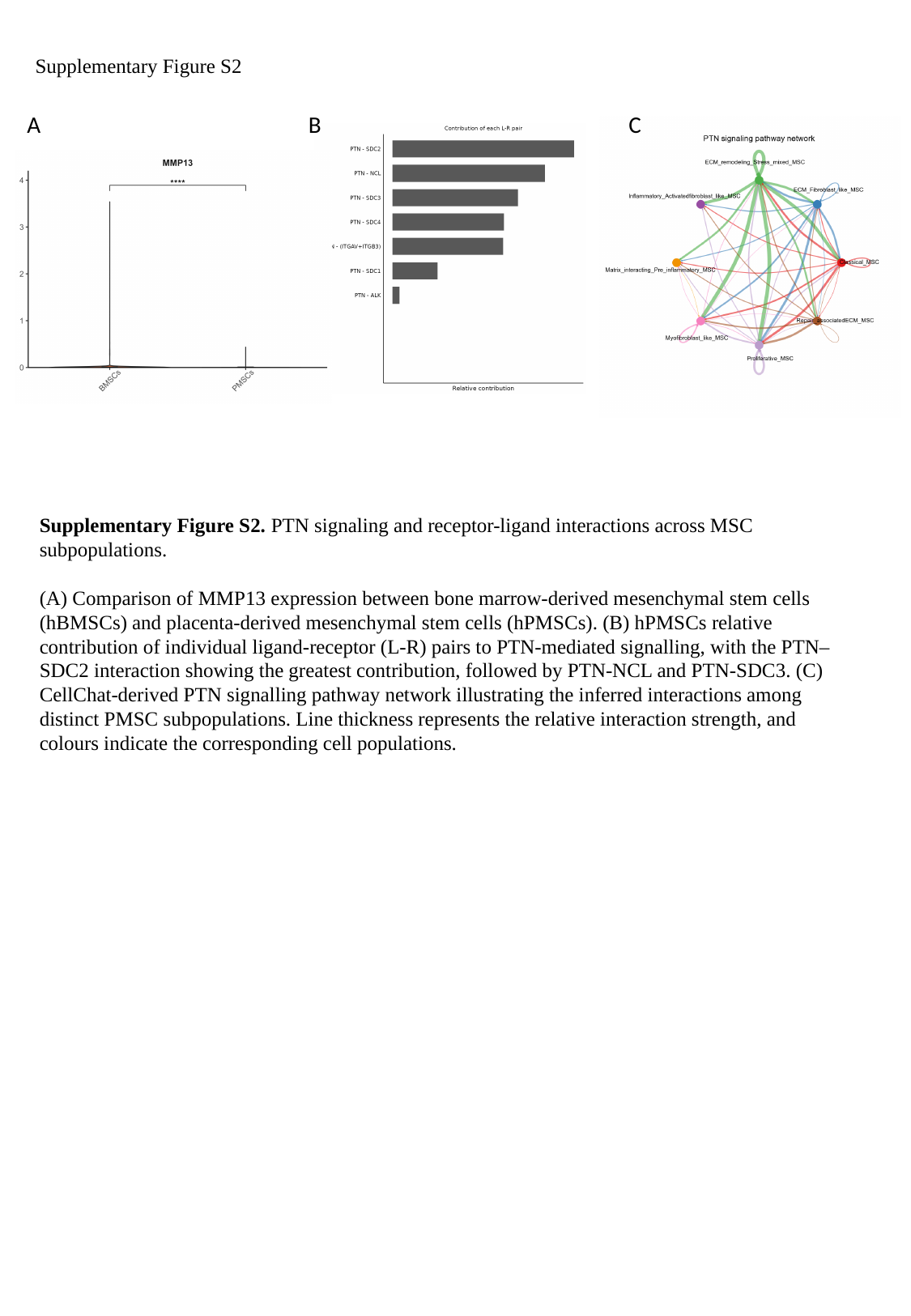

Supplementary Figure S2
A
B
C
Supplementary Figure S2. PTN signaling and receptor-ligand interactions across MSC subpopulations.
(A) Comparison of MMP13 expression between bone marrow-derived mesenchymal stem cells (hBMSCs) and placenta-derived mesenchymal stem cells (hPMSCs). (B) hPMSCs relative contribution of individual ligand-receptor (L-R) pairs to PTN-mediated signalling, with the PTN–SDC2 interaction showing the greatest contribution, followed by PTN-NCL and PTN-SDC3. (C) CellChat-derived PTN signalling pathway network illustrating the inferred interactions among distinct PMSC subpopulations. Line thickness represents the relative interaction strength, and colours indicate the corresponding cell populations.

### Slide 4
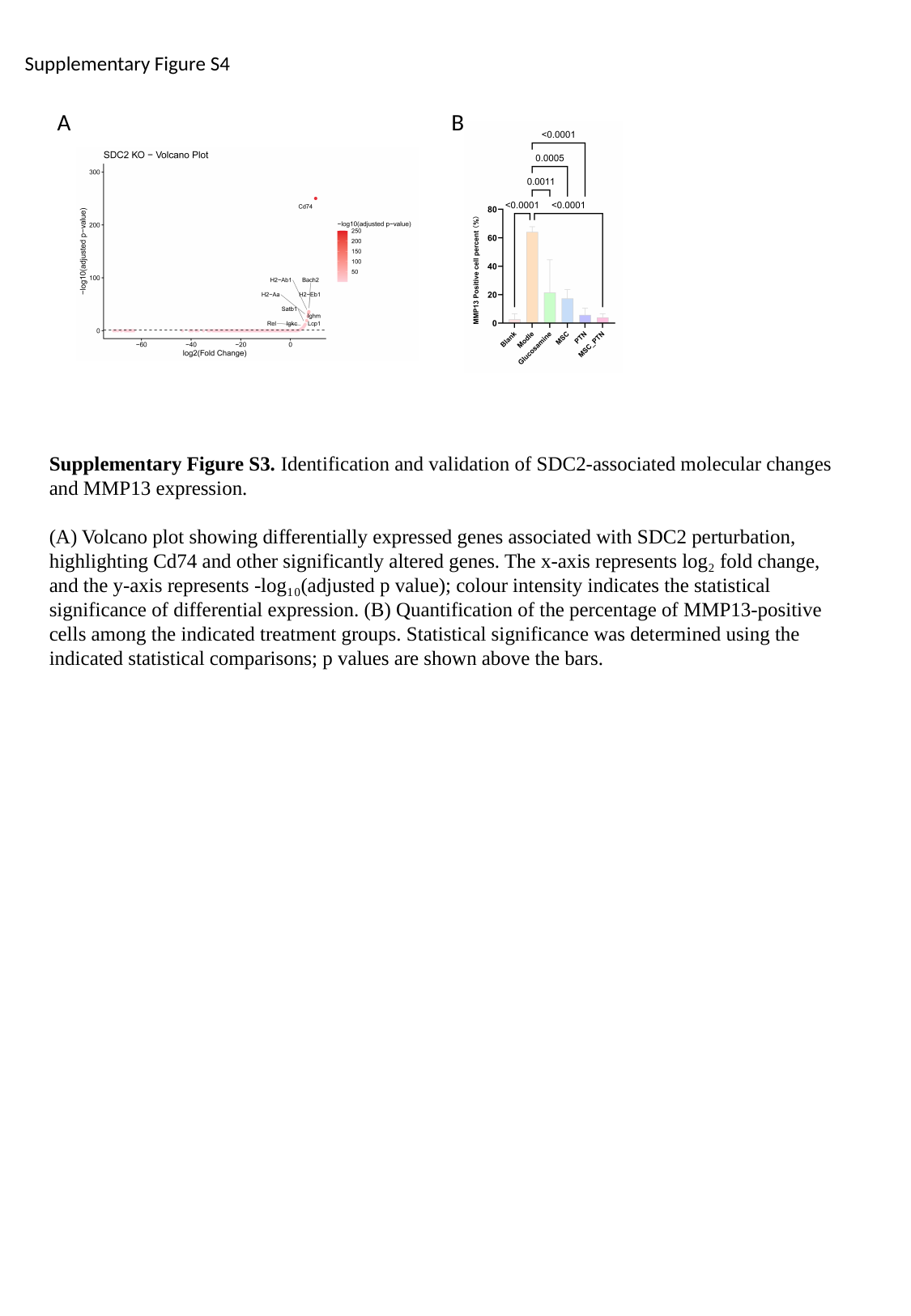

Supplementary Figure S4
A
B
Supplementary Figure S3. Identification and validation of SDC2-associated molecular changes and MMP13 expression.
(A) Volcano plot showing differentially expressed genes associated with SDC2 perturbation, highlighting Cd74 and other significantly altered genes. The x-axis represents log₂ fold change, and the y-axis represents -log₁₀(adjusted p value); colour intensity indicates the statistical significance of differential expression. (B) Quantification of the percentage of MMP13-positive cells among the indicated treatment groups. Statistical significance was determined using the indicated statistical comparisons; p values are shown above the bars.

### Slide 5
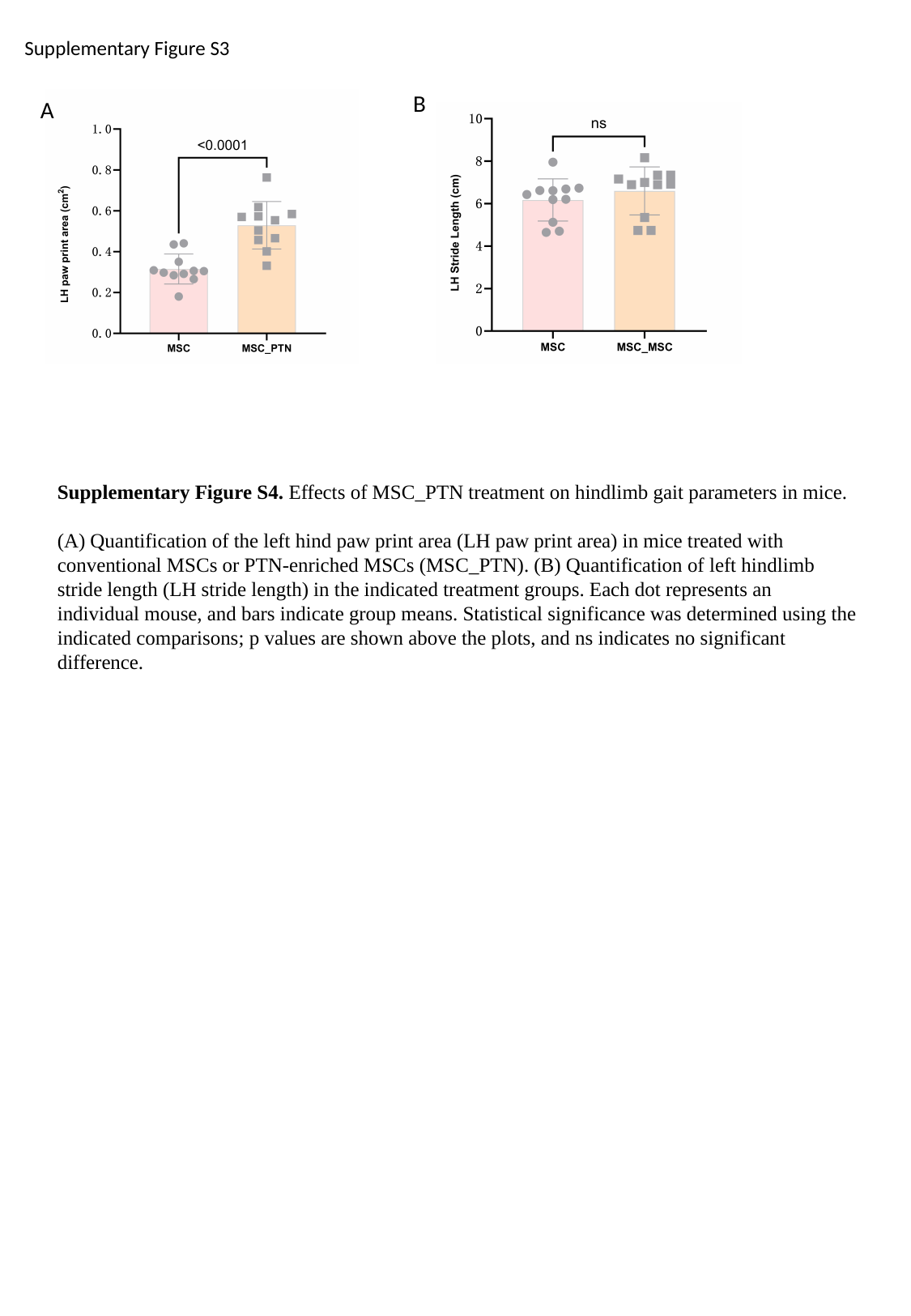

Supplementary Figure S3
B
A
Supplementary Figure S4. Effects of MSC_PTN treatment on hindlimb gait parameters in mice.
(A) Quantification of the left hind paw print area (LH paw print area) in mice treated with conventional MSCs or PTN-enriched MSCs (MSC_PTN). (B) Quantification of left hindlimb stride length (LH stride length) in the indicated treatment groups. Each dot represents an individual mouse, and bars indicate group means. Statistical significance was determined using the indicated comparisons; p values are shown above the plots, and ns indicates no significant difference.
