## Supplemental Document S3: BioRender publication license for "A PTN-Rich MSC Subtype Promotes Cartilage Repair in End-Stage Knee Osteoarthritis": Publication License 2.pdf

49 Spadina Ave. Suite 200
Toronto ON M5V 2J1 Canada
www.biorender.com

### Confirmation of Publication and Licensing Rights - Open Access

**September 9th, 2026**

| | |
|---|---|
| ***Subscription Type:*** | *Individual - Academic* |
| ***Agreement number:*** | *LA2A6LLGEG* |
| ***Publisher Name:*** | *Annals of the Rheumatic Diseases* |
| ***Figure Title:*** | *F2 flowchart* |
| ***Citation to Use:*** | *Created in BioRender. Zhao, Y. (2026) https://BioRender.com/tepb6vg* |

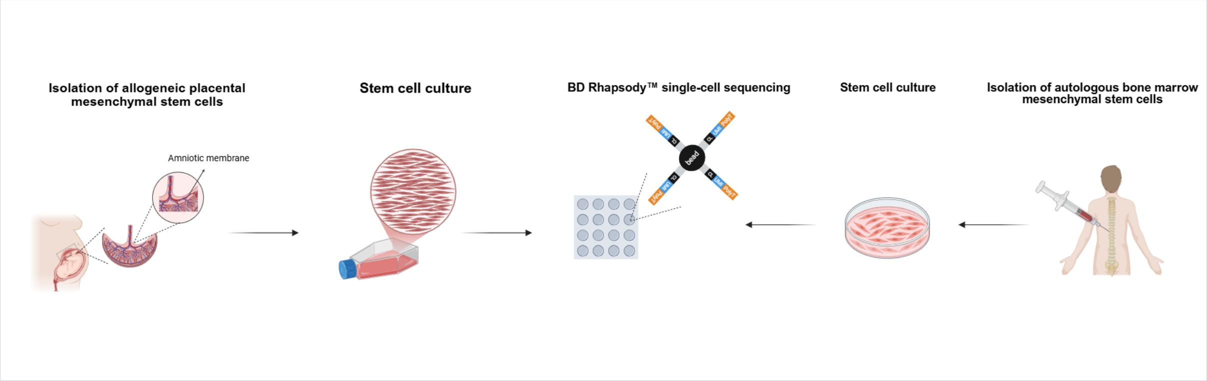


*For any questions regarding this document, or other questions about publishing with BioRender, please refer to our BioRender Publication Guide, or contact BioRender Support at.*
