## Supplementary material for "A PTN-Rich MSC Subtype Promotes Cartilage Repair in End-Stage Knee Osteoarthritis": Methods

### Sample Collection and Dissociation

### Tissue specimens (cartilage, synovium, synovial fluid) were harvested and immediately processed for single-cell profiling. Solid tissues were minced and subjected to enzymatic dissociation to generate single-cell suspensions, while synovial fluid was processed to isolate infiltrating cells. Cellular debris and dead cells were removed via density gradient centrifugation. The resulting resulted viable cells were resuspended in cold PBS supplemented with 0.04% BSA, filtered through a 40 μm cell strainer, and maintained on ice until capture.

### Single-Cell Library Preparation and Sequencing

### Cell viability and concentration were assessed using an automated cell counter, and samples with >80% viability were used for library preparation. Single-cell libraries were generated using the BD Rhapsody™ Single-Cell Analysis System, BD Rhapsody™ cDNA Kit, and BD Rhapsody™ WTA Amplification Kit according to the manufacturer’s instructions, followed by sequencing on the Illumina NovaSeq 6000 platform.

### Bioinformatics Analysis

Raw sequencing data were processed using the BD Rhapsody™ Analysis Pipeline. Quality control, normalization, dimensionality reduction (PCA and UMAP), clustering, and differential expression analysis were performed using Seurat v4.0. Cell-cell communication was inferred using CellChat, and pathway enrichment analysis was performed using clusterProfiler.

### Chromium 10x Genomics library and sequencing

Single-cell suspensions were processed using the 10x Genomics Chromium Single-Cell 3′ Kit (v3) according to the manufacturer’s instructions. cDNA amplification and library preparation were performed following the standard protocol, and libraries were sequenced on the NovaSeq X (Illumina).

**Immunofluorescence staining of cultured cells**

HEK-293T cells were seeded in six-well plates and cultured to approximately 70% confluence. Cells were transfected with PTN-mCherry alone or co-transfected with PTN-mCherry and NCL-eGFP or SDC2-eGFP. Briefly, plasmid DNA was diluted in Opti-MEM and mixed with the corresponding transfection reagent, followed by incubation for 10-20 min before addition to the cells. After 24 h of transfection, cells were fixed with 4% paraformaldehyde for 15 min at room temperature. For intracellular antigen detection, cells were permeabilised with 1% Triton X-100 in PBS for 30 min and washed three times with PBS. Non-specific binding was blocked with 5% BSA for 1 h at room temperature. Nuclei were counterstained with DAPI for 5 min in the dark, followed by three washes with PBS. Fluorescence signals from PTN-mCherry, NCL-eGFP, SDC2-eGFP and DAPI were visualised using a confocal microscope. Images were acquired at appropriate magnifications to assess the subcellular distribution and co-localisation of the fluorescent signals.

**Co-immunoprecipitation assay**

Co-immunoprecipitation was performed to validate the association of PTN with candidate receptors. HEK-293T cells were co-transfected with HA-PTN-mCherry and Flag-NCL-eGFP or Flag-SDC2-eGFP. Cells co-transfected with HA-PTN-mCherry and Flag-EGFP served as a negative control. After 36-48 h, cells were lysed in NP-40 lysis buffer containing protease inhibitors, and lysates were clarified by centrifugation at 12,000 ×g for 10-15 min at 4°C. Following retention of 5-10% of the lysate as input, the remaining lysates were incubated with anti-FLAG M2 affinity gel overnight at 4°C. After extensive washing, immunoprecipitated proteins were eluted in 2× SDS-PAGE sample buffer and analysed by western blotting using anti-HA and anti-FLAG antibodies. HA-PTN enrichment in the Flag-immunoprecipitated fraction relative to the Flag-EGFP control was used to assess the association between PTN and NCL or SDC2.

### Isolation and Culture of Mesenchymal Stem Cells

Human placental mesenchymal stem cells (hPMSCs) were isolated from the amniotic membrane of placentas obtained from healthy full-term pregnancies. Placental tissues were collected after written informed consent was obtained from all donors, and all procedures were approved by the Institutional Ethics Committee of the Affiliated Hospital for Neck, Shoulder, Waist and Leg Pain, Shandong First Medical University, in accordance with the Declaration of Helsinki (Ethics Review Approval No. 202401045).

Briefly, placental tissues were processed under sterile conditions. After separation of the amniotic membrane, the tissue was minced into fragments of approximately 1 mm³ and explanted into tissue culture flasks for primary culture. MSCs were maintained in phenol red‑free MSC medium (Tongli Haiyuan, China; Cat# AS‑33) at 37 °C in a humidified incubator with 5% CO₂. The culture medium was replaced every 2-3 days. Cells were passaged at 80-90% confluence using 0.25% trypsin‑EDTA at a split ratio of 1:3. Cells from passages 3-5 (P3-P5) were used for all subsequent experiments. The mesenchymal phenotype of cultured MSCs was confirmed by flow cytometry according to the minimal criteria established by the International Society for Cell and Gene Therapy (ISCT).

Human bone marrow-derived mesenchymal stem cells (hBMSCs) were isolated from bone marrow aspirates collected after written informed consent under the same ethical approval (Ethics Review Approval No. 202401045). Briefly, bone marrow aspirates were diluted with PBS and subjected to density-gradient centrifugation using Ficoll-Paque PLUS [manufacturer, country; Cat. No.]. The mononuclear cell fraction was collected, washed with PBS, and seeded into tissue culture flasks in [culture medium, manufacturer, country; Cat. No.]. Cells were maintained at 37 °C in a humidified incubator with 5% CO₂, with the culture medium replaced every 2-3 days. At 80-90% confluence, cells were detached with 0.25% trypsin-EDTA and passaged at a split ratio of 1:3. Cells from passages 3-5 (P3-P5) were used for subsequent experiments. The mesenchymal phenotype of hBMSCs was confirmed by flow cytometry according to the minimal criteria established by the International Society for Cell and Gene Therapy (ISCT).

### Flow Cytometric Characterization and Cell Sorting

Cultured MSCs were characterized by flow cytometry according to the minimal immunophenotypic criteria defined by the ISCT. Cells were positive for the mesenchymal stem cell markers CD73 (344015, BioLegend), CD90 (328109, BioLegend), and CD105 (323207, BioLegend), and negative for the hematopoietic markers CD34, CD45, and HLA‑DR.

For cell sorting, MSCs were dissociated into single‑cell suspensions and incubated with fluorochrome‑conjugated antibodies against CD73, CD90, and CD105 (BioLegend, USA). Cells co‑expressing high levels of CD73, CD90, and CD105 (CD73⁺CD90⁺CD105⁺) were isolated by fluorescence‑activated cell sorting (FACS) and used for all subsequent experiments.

**Animal model of knee osteoarthritis**

All animal experiments were approved by the Medical Ethics Committee of the Affiliated Hospital for Neck, Shoulder, Waist and Leg Pain, Shandong First Medical University (approval No.202502027). All animal experiments were approved by the Institutional Animal Care and Use Committee and conducted in accordance with established animal welfare guidelines.

Eight‑week‑old male C57BL/6J mice (20-25g; Jiangsu Huachuang Xinnuo Pharmaceutical Technology Co., Ltd., China) were acclimatised for one week prior to surgery. Knee osteoarthritis (KOA) was induced by bilateral anterior cruciate ligament transection (ACLT). Mice were anesthetized by intraperitoneal injection of 1.25% Avertin (2,2,2-tribromoethanol) before subsequent experimental procedures. Under general anaesthesia and sterile conditions, a medial parapatellar approach was used to expose the knee joint, followed by complete transection of the ACL under microscopic guidance. Joint instability was confirmed by the anterior drawer test. The wound was closed with 5‑0 absorbable sutures, with the same procedure performed on the contralateral knee. For postoperative care, mice were placed in a warm recovery cage with a heating pad. Knee joints were harvested eight weeks after surgery, and articular cartilage damage was evaluated using the knee Osteoarthritis Research Society International (OARSI) scoring system.

**Assessment of joint function by CatWalk gait analysis**

To evaluate the functional recovery of KOA-associated joint impairment following treatment with PTN-rich CD74-low MSCs, hindlimb locomotor performance was assessed using the CatWalk XT automated gait analysis system (Noldus Information Technology, Wageningen, The Netherlands). Before behavioural assessment, mice were acclimatised to the experimental environment for 30 min and trained for 2-3 consecutive days to ensure stable and voluntary traversal of the walkway.

During data acquisition, mice were allowed to ambulate freely across the illuminated glass walkway. Only runs with continuous, uninterrupted locomotion and meeting predefined quality criteria (run duration between 0.5 and 5.0 s and maximum speed variation ≤60%) were included for subsequent analysis. At least three valid crossings were recorded for each animal at each time point (35 days after MSC injection). Gait parameters were automatically extracted using CatWalk XT software, including stride length, paw print area, and paw print intensity, which were used as indicators of functional recovery following MSC‑based intervention.

**Histological analysis and immunofluorescence**

Knee joints were dissected and decalcified in 10% EDTA for two weeks prior to paraffin embedding for histological evaluation, including Safranin O/Fast Green (SO/FG) staining, haematoxylin and eosin (H&E) staining, and immunofluorescence (IF) Following complete decalcification, specimens were dehydrated through a graded ethanol series, cleared in xylene, embedded in paraffin, and serially sectioned in the sagittal plane at a thickness of 5 μm.

For histological assessment of cartilage repair, paraffin sections were deparaffinised, rehydrated, and subjected to H&E staining to evaluate overall joint morphology and tissue organisation, as well as SO/FG staining to assess cartilage matrix integrity and proteoglycan content. Cartilage degeneration was evaluated using the knee OARSI scoring system, as described above.

For immunofluorescence analysis, sections were deparaffinised and rehydrated, followed by heat-induced antigen retrieval in citrate buffer (pH 6.0) at 95-100°C for 15 min. After permeabilisation with 0.3% Triton X-100 and blocking with goat serum, sections were incubated overnight at 4°C with primary antibodies against SOX9 (ab185966, Abcam, 1:200) , MMP13 (ab39012, Abcam, 1:200) and IL-6 (GB11117, Servicebio, 1:200). Following washes, sections were incubated with fluorophore‑conjugated secondary antibodies (Cy3 conjugated Goat Anti-Rabbit IgG (H+L)，GB27303，Servicebio, 1:400，Cy3550/570) for 1h at room temperature in the dark. Nuclei were counterstained with DAPI (1 μg/mL), and sections were mounted with antifade mounting medium. Images were acquired using a confocal laser scanning microscope under identical acquisition parameters for all samples to enable quantitative comparison.

**Protein extraction and Western blot**

Bone samples were pulverised under liquid nitrogen following removal of soft tissues and bone marrow. Total protein was extracted in pre-chilled RIPA buffer supplemented with protease and phosphatase inhibitors and quantified using a BCA assay. Equal amounts of protein (50 μg) were separated by SDS-PAGE, transferred to PVDF membranes, and immunoblotted with anti-CD74 antibody (Abcam, ab289885, 1:1000) overnight at 4°C, followed by Goat Anti RabbitlgG (H+L)-HRP secondary antibody (Diagbio, db10002, 1:5000). Signals were detected using ECL and imaged using a gel documentation system.

***In vitro* indirect co-culture assay**

To establish a non‑contact co‑culture system, Transwell (Corning, 723101) inserts with a pore size of 0.4  μm were employed. Prior to the experiment, cartilage (CA), synovium (SY), and synovial fluid (SF), as well as PTN-rich CD74‑low MSCs, were isolated and cultured separately. When all cell types reached the logarithmic growth phase, with viability ≥ 95% and negative results in sterility testing, they were harvested by trypsin (Gibco, 25200072) digestion, centrifuged, and resuspended to prepare single‑cell suspensions. The Transwell inserts and culture plates were pre‑sterilised, and the culture medium as well as PBS were pre‑warmed for subsequent use.

SY, CA, and SF cells were seeded into the lower compartment of the Transwell system at a density of 1 × 10⁴ cells per well and incubated under standard conditions (37 °C, 5% CO₂, saturated humidity) for 4 h to allow complete cell attachment. Subsequently, the Transwell inserts were placed into the wells, and PTN-rich CD74‑low MSCs were seeded into the upper compartment at a density of 5 × 10³ cells per well. After confirming the absence of leakage, the co‑culture was continued for 48  h under the same incubation conditions without medium replacement during this period. At 24 h and 48 h, cell growth status and system integrity were examined under an inverted microscope to ensure the absence of contamination, leakage, or abnormal apoptosis.

Upon completion of the co‑culture at 48 h, the upper inserts and the spent medium were carefully removed under aseptic conditions, and the culture medium in the lower compartment was discarded. The cells in the lower compartment were then gently digested with 0.25% trypsin‑EDTA at 37 °C, and the enzymatic reaction was terminated by adding complete culture medium. The resulting cell suspension was collected and centrifuged at 1,000 rpm for 5 min to obtain cell pellets. The pellets were washed by resuspension in PBS to remove residual impurities, ultimately yielding highly purified target cells from the lower compartment for subsequent analyses.

**RNA extraction and qPCR**

Mouse femur/tibia was snap-frozen in liquid nitrogen and ground to a fine powder. Approximately 50 mg of tissue powder was lysed in 500 μL FreeZol Reagent (Vazyme, R711-01), incubated at room temperature for 5 min, and centrifuged at 12,000×g for 5 min. The supernatant was mixed with 100 μL Dilution Buffer, incubated for 5 min, and centrifuged at 12,000×g for 15 min. The upper aqueous phase was collected and mixed with an equal volume of isopropanol to precipitate RNA. After centrifugation at 12,000×g for 10 min, the pellet was washed twice with 75% ethanol (8,000×g, 3 min), air-dried, and dissolved in RNase-free water. One microgram of total RNA was treated with 4× gDNA wiper Mix (Vazyme, R323-01) at 42°C for 2 min to remove genomic DNA, followed by reverse transcription using 5× HiScript III qRT SuperMix (total 20 μL). Quantitative PCR was performed in a 20 μL reaction containing 2× TaqPro Universal SYBR Master Mix (Vazyme, Q711-02), 0.2 μM primers, and cDNA template, using a standard thermal cycling protocol on a real-time PCR system.The primer sequences used in this study are provided in Supplementary Table S2.

**Statistical analysis**

Statistical analysis was performed using GraphPad Prism 11.0 (GraphPad Software, San Diego, CA, USA). All data are presented as mean ± standard deviation (SD). For comparisons among multiple independent groups, one‑way analysis of variance (ANOVA) was applied, followed by Dunnett's multiple comparisons test for pairwise comparisons between control and experiment groups. A value of *p* < 0.05 was considered statistically significant (**P* < 0.05, ***P* < 0.01, ****P* < 0.001, and *****P* < 0.0001).

**Scientific illustration**

Scientific schematics and graphical illustrations were generated using BioRender (BioRender.com) under an active publication licence (Supplementary Document S3).
